# Non-Canonical Activation of CREB Mediates Neuroprotection in a *C. elegans* Model of Excitotoxic Necrosis

**DOI:** 10.1101/261420

**Authors:** K. Genevieve Feldmann, Ayesha Chowdhury, Jessi Becker, N’Gina McAlpin, Taqwa Ahmed, Syed Haider, Jian X. Richard Xia, Karina Diaz, Monal G. Mehta, Itzhak Mano

**Affiliations:** Department of Molecular, Cellular and Biomedical Sciences, CDI Cluster on Neural Development and Repair, The CUNY School of Medicine, City College (CCNY), The City University of New York (CUNY); The CUNY Neuroscience Collaborative PhD Program, CUNY Graduate Center; Undergraduate Program in Biology, CCNY, CUNY; The Sophie Davis BS/MD program, CUNY School of Medicine; Robert Wood Johnson Medical School, Rutgers – The State University of New Jersey

**Keywords:** C. elegans, CREB, CRTC, Excitotoxicity, Neurodegeneration, Neuroprotection

## Abstract

Excitotoxicity, caused by exaggerated neuronal stimulation by Glutamate (Glu), is a major cause of neurodegeneration in brain ischemia. While we know that neurodegeneration is triggered by overstimulation of Glu-Receptors (GluRs), the subsequent mechanisms that lead to cellular demise remain controversial. Surprisingly, signaling downstream of GluRs can also activate neuroprotective pathways. The strongest evidence involves activation of the transcription factor cAMP Response Element Binding-protein (CREB), widely recognized for its importance in synaptic plasticity. Canonical views describe CREB as a phosphorylation-triggered transcription factor, where transcriptional activation involves CREB phosphorylation and association with CREB Binding Protein (CBP). However, given CREB’s ubiquitous cross-tissue expression, the multitude of cascades leading to CREB phosphorylation, and its ability to regulate thousands of genes, it remains unclear how CREB exerts closely-tailored, differential neuroprotective responses in excitotoxicity. A non-canonical, alternative cascade for activation of CREB-mediated transcription involves the CREB co-factor cAMP-regulated transcriptional co-activator (CRTC), and may be independent of CREB phosphorylation. To identify cascades that activate CREB in excitotoxicity we use a *C. elegans* model of neurodegeneration by excitotoxic necrosis. We demonstrate that CREB’s neuroprotective effect is conserved, and seems most effective in neurons with moderate Glu exposure. We find that factors mediating canonical CREB activation are not involved. Instead, phosphorylation-independent CREB activation in nematode excitotoxic necrosis hinges on CRTC. CREB-mediated transcription that depends on CRTC, but not on CREB phosphorylation, might lead to expression of a specific subset of neuroprotective genes. Elucidating conserved mechanisms of excitotoxicity-specific CREB activation can help us focus on core neuroprotective programs in excitotoxicity.

## Introduction

Excitotoxic neurodegeneration is a leading cause of neuronal damage in brain ischemia, and a contributing factor in a range of progressive neurological diseases (Choi & Rothman 1990, Moskowitz *et al.* 2010, Lai *et al.* 2014, Tymianski 2014). Excitotoxicity is trigged by malfunction of Glutamate Transporters (GluTs) (Danbolt 2001), leading to the accumulation of Glutamate (Glu) in the synapse, overstimulation of Glu Receptors (GluRs) on postsynaptic neurons, and postsynaptic buildup of toxic Ca^2+^ concentrations. The exaggerated Ca^2+^ influx eventually causes extensive neurodegeneration, with morphology that spans the range from necrosis to apoptosis, correlated with the extent of the insult (Choi & Rothman 1990, Mehta *et al.* 2007, Moskowitz et al. 2010, Lai et al. 2014). Neuronal destruction in brain ischemia is progressive: Neuronal damage is most rapid and severe in the core of the stroke area, while the surrounding penumbra is initially only “stunned”. The fate of neurons in the penumbra (degeneration by necrosis, by apoptosis, or recovery) becomes apparent only at a later stage. The neurons in the penumbra area are therefore considered salvageable if appropriate therapy can be devised to protect them (Moskowitz et al. 2010). Although a plethora of intricate signaling cascades has been proposed to mediate the toxic effect of GluR hyperactivation (Mehta et al. 2007, Lai *et al.* 2011, Tymianski 2011, Fan *et al.* 2017), the contribution of each of these mechanisms to excitotoxicity *in vivo* is fiercely debated. Furthermore, a large number of clinical trials based on GluR antagonists or inhibitors of the proposed immediate downstream mechanisms have failed (Ikonomidou & Turski 2002, Tymianski 2014, Lai et al. 2014). These failures suggested that in the clinical setting, application of GluR antagonists “missed the boat”, as neurodestructive processes are already well underway by the time treatment is administered (Ikonomidou & Turski 2002)

However, these failures also emphasized a novel perspective: Surprisingly, in addition to its role in neurodegeneration, GluR activation also triggers neuroprotective signaling cascades that can mitigate neuronal demise. These neuroprotective effects were initially studied when protection was triggered at the same time of-, or even before-hyperstimulation of GluR (the latter being an example of preconditioning) (Meller *et al.* 2005, Gidday 2006, Hardingham & Bading 2010, Kitagawa 2012, Lai et al. 2014). In both cases, the transcription factor cAMP Response Element Binding protein (CREB) has a central role (Walton & Dragunow 2000, Ikonomidou & Turski 2002, Sakamoto *et al.* 2011, Lai *et al.* 2014). Importantly, subsequent studies showed that activating GluR- and CREB-mediated neuroprotection has beneficial effects even if triggered considerably after the onset of the excitotoxic insult, marking it with special clinical relevance (Papadia *et al.* 2005, Liu *et al.* 2007, Zhao *et al.* 2012).

CREB was originally identified as a bZIP-domain containing transcription factor (TF) that is widely found in many cell types, and is activated by phosphorylation on Ser133 (a residue located within a kinase-inducible transactivation domain, or KID domain) by the cAMP-dependent kinase PK-A (Montminy & Bilezikjian 1987, Yamamoto *et al.* 1988, Gonzalez & Montminy 1989, Goodman 1990). With time, CREB was marked as a prototypic example of a TF activated by phosphorylation by a range of kinases (Mayr & Montminy 2001, West *et al.* 2001, Lonze & Ginty 2002, Deisseroth *et al.* 2003). CREB was further identified as a central TF in neuroscience, as it was found to mediate the changes in gene expression that occur in synaptic plasticity (Dash *et al.* 1990, Sheng *et al.* 1990, Kandel 2001, West *et al.* 2002, Deisseroth et al. 2003, Carlezon *et al.* 2005), a function that is conserved throughout evolution (Yin *et al.* 1995, Kandel 2001). Ca^2+^ was determined to be the central trigger of CREB phosphorylation and activation following synaptic activity (Sheng et al. 1990, Sheng *et al.* 1991, Deisseroth *et al.* 1996, Greer & Greenberg 2008), and a Histone Acetyl Transferase (HAT) protein called CREB Binding Protein (CBP) was found to bind the KID transactivation domain of phosphorylated CREB and mediate transcriptional activation (Chrivia *et al.* 1993). In the canonical view of CREB activation by synaptic activity (Figure 1A), Ca^2+^ influx through GluRs triggers the activation of Ca^2+^-Calmodulin (CaM)-dependent kinases that lead to phosphorylation of CREB on its transactivation domain, allowing it to recruit CBP and stimulate transcription. Ca^2+^-induced CREB phosphorylation is suggested to come from the moderate activation of the CREB-kinase CaMK-IV by CaM in the nucleus, an effect that is strongly augmented with the robust activation of CaMK-IV by the CaMK-IV-activating kinase CaMKK (Impey & Goodman 2001, Lonze & Ginty 2002, West et al. 2002, Wayman *et al.* 2008, Flavell & Greenberg 2008). Some studies suggest that there is special importance to a nuclear Ca^2+^ signal as the critical trigger for CaMK-IV activation (Bading 2013). The same mechanisms were also suggested to be at work to provide neuroprotection from excitotoxicity (Walton *et al.* 1999, Walton & Dragunow 2000, Mabuchi *et al.* 2001, Mantamadiotis *et al.* 2002, Hardingham *et al.* 2002, Ikonomidou & Turski 2002, Kitagawa 2007, Hardingham & Bading 2010, Sakamoto et al. 2011, Lai et al. 2014). Further studies identified CREB-targeted neuroprotective genes that are turned-on under conditions of canonical CREB activation, focusing on anti-apoptotic genes (Hardingham et al. 2002, Zhang *et al.* 2007, Zhang *et al.* 2009). However the target genes and mechanisms of CREB-mediated protection from excitotoxic *necrosis* remain under-explored.

**Figure 1:**
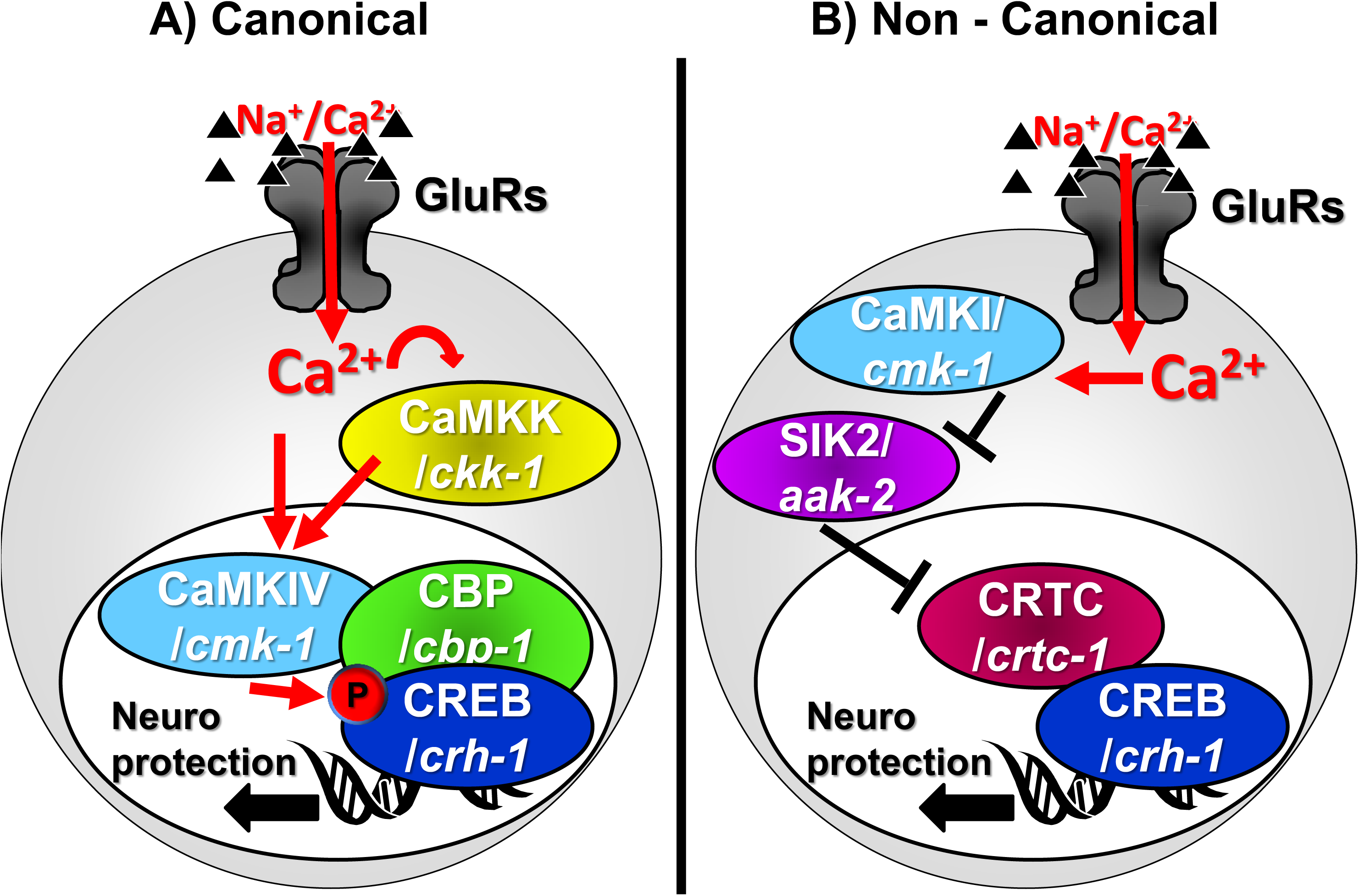
Models of CREB activation by GluR activity leading to neuroprotection. A) Canonical. B) Non Canonical.

Moreover, it is unclear how the widely-used canonical mode of CREB activation is able to specify a closely-tailored pattern of gene expression specific for excitotoxic neuroprotection. Indeed CREB is ubiquitously expressed in all tissues (Mayr & Montminy 2001), it can be stimulated by hundreds of different stimuli (Johannessen *et al.* 2004), and it regulates the expression of a total of 4,000-6,000 different genes (Zhang *et al.* 2005, Impey *et al.* 2004). Only a subset of these genes is suggested to be expressed in any given scenario (Cha-Molstad *et al.* 2004, Zhang et al. 2005). Further specificity in CREB’s action is believed to be conferred by the constellation of cellular conditions that trigger CREB activation in each case. This variability in activation modes might in turn give rise to diversification in the way CREB is activated and its interaction with other TFs (Lyons & West 2011). For instance, a subset of CREB triggering conditions leads to the recruitment members of the family of CREB co-factors called cAMP-regulated transcriptional co-activators (CRTCs), one of which (CRTC1) is specifically expressed in the brain (Wu *et al.* 2006, Altarejos *et al.* 2008, Watts *et al.* 2011). Unlike CBP, CRTCs bind directly to CREB’s bZIP DNA-binding domain, to stabilize CREB’s interaction with the DNA (Iourgenko *et al.* 2003, Conkright *et al.* 2003, Screaton *et al.* 2004, Takemori *et al.* 2007, Altarejos & Montminy 2011). CRTC includes its own transactivation domain, adding to the capability of the complex to bind additional factors and activate transcription (Altarejos & Montminy 2011). The availability of CRTC in the nucleus depends on its own phosphorylation state (Figure 1B): When CRTC is unphosphorylated, it translocates into the nucleus and activates CREB. However, under resting conditions CRTC is prevented from entering the nucleus due to its phosphorylation by the Salt Induced Kinases (SIK1/2). In the non-canonical mode of CREB activation, Ca^2+^-triggered inhibition of SIK1/2 allows CRTC to enter the nucleus and activate CREB-mediated transcription, independently of CREB phosphorylation and CBP binding. Under these conditions the CRTC::CREB complex associates instead with other HATs such as PCAF/KAT2 or KAT5 (Ravnskjaer *et al.* 2013, Clark *et al.* 2015) that mediate a different profile of histone acetylation (compared to pCREB::CBP) (Uchida *et al.* 2017). Such non-canonical, CRTC-mediated activation of CREB-mediated transcription was observed in neurons under some conditions of synaptic plasticity (Kovacs *et al.* 2007, Ch'ng *et al.* 2012, Nonaka *et al.* 2014, Briand *et al.* 2015, Uchida et al. 2017, Parra-Damas *et al.* 2017) and in Huntington’s disease (Jeong *et al.* 2012). One study further suggests that the non-canonical mode of CREB activation is also operational in excitotoxic neuroprotection, where SIK2 inhibition is achieved by Ca^2+^-mediated activation of cytoplasmic CaMK-I, which can phosphorylate and inhibit SIK2 (Sasaki *et al.* 2011). Therefore, while the prevalent view of CREB-mediated transcription in neuronal plasticity and neuroprotection is based on the canonical activation of transcription (by phosphorylation of CREB and its association with CBP), a few lines of evidence suggest instead that neuroprotection and some types of memory paradigms might be achieved by a non-canonical mode of activation. This non-canonical mechanism depends on the function of CaMK-I, SIK2, and CRTC, it is independent of CBP or CREB phosphorylation, and it might result in a different profile of histone acetylation (Altarejos & Montminy 2011, Sasaki et al. 2011, Ch'ng et al. 2012, Uchida et al. 2017). Given the paucity of evidence in support of non-canonical activation of CREB in neuroprotection, it remains unclear if this mode of CREB activation is widely involved in reducing excitotoxic damage, and if it protects from apoptosis or necrosis.

These two mechanisms of CREB activation might contribute to different modes of neuroprotection, depending on the exact conditions (such as the presence of apoptosis vs necrosis). However, while transcriptional programs for GluR-mediated protection from apoptosis have been studied before (Zhang et al. 2007, Zhang et al. 2009), evolutionarily conserved processes that promote neuronal survival in excitotoxic *necrosis* are understudied. Filling this gap might highlight the critical core of the neuroprotective pathway in this devastating form of neurodegeneration in excitotoxicity. We therefore turned to study this question in our model of excitotoxic necrosis in the nematode *C. elegans* (Mano & Driscoll 2009). In addition to the strong conservation of key cellular mechanisms (including cell death) (Lettre & Hengartner 2006), the powerful genetic (Brenner 1974) and neuroscience (White *et al.* 1986, Rankin 2002) tools available in this system make the elucidation of critical signaling cascades especially productive. Glutamate is a central neurotransmitter in the worm and is widely used to stimulate command interneurons through conserved GluRs (Brockie & Maricq 2006). GluRs mediate both basic signaling and synaptic plasticity (Rose & Rankin 2006, Rose *et al.* 2005, Rose *et al.* 2003, Emtage *et al.* 2009, Stetak *et al.* 2009). CREB (worm homolog name: CRH-1) and the components of its canonical activation cascade are also well conserved in the worm, and regulate learning and memory and synaptic plasticity (Kimura *et al.* 2002, Bates *et al.* 2006, Suo *et al.* 2006, Kauffman *et al.* 2010, Timbers & Rankin 2011, Nishida *et al.* 2011, Yu *et al.* 2014, Lakhina *et al.* 2015, Chen *et al.* 2016, Moss *et al.* 2016, Freytag *et al.* 2017, Nishijima & Maruyama 2017). CREB/CRH-1 can be activated by phosphorylation (on a site homologous to Ser133) by the nematode’s combined CaMK I/IV homolog CMK-1, whose basal activity is greatly stimulated by CaMKK/CKK-1 (Kimura et al. 2002, Yu et al. 2014). Similarly, CBP/CBP-1 cooperates with CREB/CRH-1 in many cells (Eastburn & Han 2005), and protects neurons from polyglutamine-induced neurodegeneration (Bates et al. 2006). SIK1/2 homologs are also present, including KIN-29 and AAK-2 (Lanjuin & Sengupta 2002, Savage-Dunn *et al.* 2003, Singaravelu *et al.* 2007, Apfeld *et al.* 2004, Mair *et al.* 2011). Similarly to SIK1/2 in mammals, AAK-2 modulates the nuclear translocation of CRTC/CRTC-1 to regulate CREB/CRH-1 –mediated transcription (and here too, CRTC-1 needs to be unphosphorylated in order to be preferentially localized to the nucleus) (Mair et al. 2011, Burkewitz *et al.* 2015). In our model of nematode excitotoxicity (Mano & Driscoll 2009), knockout (KO) of the cardinal GluT gene *glt-3* (Mano *et al.* 2007) in the *nuIs5* sensitized background (Berger *et al.* 1998) causes the necrotic death of some of the neurons postsynaptic to Glu connections. This Glu-triggered neuronal necrosis is independent of canonical apoptosis (Tehrani *et al.* 2014), and it shows key features conserved in excitotoxicity, such as dependence on key Ca^2+^-permeable GluRs (Brockie & Maricq 2006), Ca^2+^ release from intracellular stores, involvement of DAPK (Del Rosario *et al.* 2015), and modulation by FoxO/DAF-16 (Tehrani et al. 2014). We therefore set out to address the controversy regarding the mechanism of CREB-mediated neuroprotection in excitotoxic necrosis, using a powerful genetic model where the most important core events in this process are likely to be conserved. In the current study we find that indeed CREB/CRH-1 has a neuroprotective role in neurons exposed to a moderate excitotoxic necrosis, and that the non-canonical mechanism of CREB activation is the one that is evolutionary conserved in neuroprotection from excitotoxic necrosis.

## Methods

### Strains

Strains were maintained at 20°C according to Brenner (Brenner 1974), and grown on MYOB agar plates (Church *et al.* 1995) seeded with OP50 (Stiernagle 2006). All the major new strains were constructed twice from independent crosses, and data was verified to be similar.

**Strains used: WT:** Bristol N2 **Excitotoxicity strain:** ZB1102: *glt-3(bz34) IV; nuIs5 V* (Mano & Driscoll 2009) ***crh-1:*** YT17: *crh-1(tz2) III* (Kimura et al. 2002) ***age-1:*** TJ1052: *age-1(hx546) II* (Friedman & Johnson 1988), ***cbp-1 hyperactivity:*** MH2430: *cbp-1(ku258) III* (Eastburn & Han 2005), ***cmk-1:*** VC220: *cmk-1(ok287) IV; gkDf56 Y102A5C.36(gk3558) V* (Satterlee *et al.* 2004), ***aak-2:*** RB754: *aak-2(ok524) X* (Apfeld et al. 2004) ***crtc-1:*** *crtc-1(tm2869) I* (Mair et al. 2011) ***rol-6:*** HE1006: *rol-6(su1006) II*, **WT CRTC labeled with RFP:** AGD418: *uthIs205[P_crtc..1_::crtc-1::RFP::unc-54 3'UTR; rol-6(su1006)]* (Mair et al. 2011) **Unphosphorylatable CRTC:** AGD466: *uthEx222*[*P_crtc-1::_crtc-1 _c_DNA(S76A, S179A)::tdTomato::unc-54 3’UTR; rol-6(su1006)]****Glutamatergic behavioral negative control:*** VM1268: *nmr-1(ak4) II; glr-2(ak10) glr-1(ky176) III.* Some strains were obtained from *C. elegans* Genetic Center (CGC), Japanese National Bioresource Project (NBRP) or from the original creators. **Strains created in this study: *crh-1 in excitotoxicity*** (by crossing YT17 and ZB1102) IMN36: *crh-1(tz2) III; glt-3(bz34) IV; nuIs5 V;* ***age-1 in excitotoxicity*** (by crossing TJ1052 and ZB1102) IMN37: *age-1(hx546) II; glt-3(bz34) IV; nuIs5 V;* ***age-1 and crh-1 epistasis in excitotoxicity*** (by crossing IMN36 and IMN37) IMN38: *age-1(hx546) II; crh-1(tz2) III; glt-3(bz34) IV; nuIs5 V;* ***cbp in excitotoxicity*** (by crossing MH2430 and ZB1102) IMN39: *cbp-1(ku528) III; glt-3(bz34) IV; nuIs5 V;* ***cmk-1 in excitotoxicity*** (by crossing VC220 and ZB1102) IMN40: *cmk-1(ok287) IV; glt-3(bz34) IV; nuIs5 V* ***aak-2 in excitotoxicity*** (by crossing RB754 and ZB1102) IMN41: *glt-3(bz34) IV; nuIs5 V; aak-2(ok524) X;* ***crtc in excitotoxicity*** (by crossing *crtc-1(tm2869) I* and ZB1102) IMN42: *crtc-1(tm2869) I; glt-3(bz34) IV; nuIs5 V;* ***crtc & aak-2 epistasis in excitotoxicity*** (by crossing IMN41 and IMN42) IMN43: *crtc-1(tm2869) I; glt-3(bz34) IV; nuIs5 V; aak-2(ok524) X;* ***roller phenotype control in excitotoxicity*** (by crossing HE1006 and ZB1102) IMN44 *rol-6(su1006) II; glt-3(bz34) IV; nuIs5 V;* **WT CRTC overexpression *in excitotoxicity*** (by crossing AGD418 and ZB1102): IMN45 glt-3(bz34) IV; nuIs5 V; uthIs205[P__crtc-1_:_*::crtc-1::RFP::unc-54 3’UTR; rol-6(su1006)];***unphosphorylatable CRTC overexpression in excitotoxicity:** (by crossing AGD466 and ZB1102) IMN46: *glt-3(bz34) IV; nuIs5 V; uthEx222[P_crtc-1_::crtc-1 cDNA (S76S, S179A)::tdTomato::unc-54 3’UTR; rol-6(su1006)];***WT CREB rescue:** IMN47 *crh-1(tz2) III; glt-3(bz34) IV; nuIs5 V; Ex[P_glr_::crh-1 cDNA::dsRed; P_mec-4_::RFP];***Phosphorylation mutant CREB rescue** IMN48: *crh-1(tz2) III; glt-3(bz34) IV; nuIs5 V; Ex[P_glr_::S29A crh-1 cDNA::dsRed; P_mec-4_::GFP]* All strains were confirmed homozygous using PCR. Transgenes expressing green or red fluorescence were followed by microscopy.

### Neurodegenerative Quantification

Neurodegeneration was quantified using an inverted scope and Nomarski Differential Interference Contrast (DIC) according to (Mano & Driscoll 2009, Tehrani et al. 2014, Del Rosario et al. 2015). These necrotic neurons appear as vacuolar-looking structures as previously described. The data was confirmed in independent lines. The data collection was blinded.

### Fluorescence Microscopy

We mounted L3 animals on agarose gel pads (2%) microscope slides with a drop of M9 buffer. 10mM sodium azide (NaN_3_) was added to immobilize worms for CRTC localization. Strains AGD418 (Mair et al. 2011) and IMN41 were imaged using the 100X objective oil lens; z-stacks were taken using the multi-dimensional acquisition tool using Metamorph Software. Images were examined using ImageJ (imagej.nih.gov) and deconvolved using Autoquant X3.

### Identification of Degenerating Neurons

L3 animals from strains ZB1102 and IMN32 were imaged with Nomarski DIC and fluorescence microscopy (AxioZeiss) with the 63X objective. Some animals were treated with 10mM NaN_3_ and showed the same degeneration pattern as animals that were analyzed without the use of NaN_3_. DIC and GFP images were examined independently and in merged format in ImageJ. To identify the degenerating neuron, the location of DIC and GFP-labeled cell body and the shape of the neuron’s processes were compared to those of glr-1–expressing neurons (Brockie & Maricq 2006, Altun *et al.* 2002-2018).

### Molecular Biology

Two pENTR Gateway vectors, one expressing wildtype CREB/*crh-1* cDNA and a the other expressing CREB/crh-1 cDNA with a single point mutation making a phosphorylation mutant (S29A) were a gift from Hidehito Kuroyanagi (Kimura et al. 2002). In addition, Vector KP#889 (*P_glr-1_::dsRed* (a gift from the Kaplan and the Juo labs) (Kowalski *et al.* 2011), was used as the destination vector. Plasmid expressing *P_mec-4_::GFP* and *P_mec-4_::mCherry* (Gifts from Driscoll lab (Royal *et al.* 2005, Toth *et al.* 2012)) were used as coinjection markers. WT *crh-1* and S29A *crh-1* cDNAs were inserted at the BamHI site of KP#889 vector to create a final vectors: *P_glr-1_::crh-1 cDNA(wt)::dsRed* and *P_glr-1_::crh-1 cDNA(S29A)::dsRed.* Final plasmids were confirmed with sequencing using a 5’ CTTCGTCTCGGTCACTTCACTTCG primer for the KP#889 promoter of *glr-1.* Plasmids were injected into IMN36 *crh-1;glt-3;nuIs5* worms (L4 or Young Adult) using standard protocols. Two independent lines were maintained for each wildtype and mutant strains.

### Behavioral assays

Locomotion assays (duration of spontaneous forward mobility) (Brockie *et al.* 2001b) and nose touch assays (NOT) (Kaplan & Horvitz 1993, Hart *et al.* 1995) assays were performed blindly.

### Statistics

Pairwise comparisons were made using Student’s two-tailed t-test with a significance threshold of *p<0.05.* SEM was also calculated. Degeneration frequency was calculated as the sum of neuron (individual or by type) divided by the total possible neurons dying. Averages and SEMs were also calculated.

## Results

### The neuroprotective role of CREB/CRH-1 is conserved in nematode excitotoxicity

We tested the effect of CREB in nematode excitotoxicity by introducing a standard knockout (KO) allele of the CREB homolog gene (*crh-1(tz2)III*, missing the bZIP domain) (Kimura et al. 2002) into our excitotoxicity strain (*glt-3(bz34) IV; nuIs5 V)* (Mano & Driscoll 2009). The nematode excitotoxicity strain combines a KO of the centrally important GluT gene *glt-3* with the sensitizing transgenic modification *nuIs5.* The *nuIs5* transgene puts ∼30 identified neurons postsynaptic to Glu connections at risk of neurodegeneration by expressing a hyperactive Gαs and GFP under the promoter of the GluR subunit *glr-1* (Berger et al. 1998). Combining the *nuIs5* sensitized background with the GluT KO mutation *glt-3(bz34)* gives rise to seemingly stochastic- (but see below), and GluR-dependent-necrosis of some of these at-risk neurons. We quantify the extent of necrosis by observing animals at different developmental stages and counting the number of swollen neurons in the head (by screening through a large number of live animals in a mixed population, identifying developmental stage and counting vacuole-like structures in each animal using DIC optics). Typically, excitotoxic neurodegeneration peaks at the L3 developmental stage (coinciding with the maturation of Glu signaling) at the level of ∼4.5 head neurons/animal (a number that later declines, probably due to removal of cell corpses by engulfment) (Mano & Driscoll 2009).

We now observe that adding the CREB/*crh-1 ko* to the excitotoxicity strain *(glt-3;nuIs5)* caused a dramatic surge in the extent of necrotic neurodegeneration, increasing the average number of dying head neurons per animal at L3 from 4.5 to above 8 (Figure 2). These results suggest that the presence of fully functional CREB/CRH-1 in wild type (wt) animals protects some of the at-risk neurons from excitotoxic necrosis, and its absence in the *crh-1 ko* strain causes more of these at-risk neurons to die. We therefore conclude that, like many other aspects of our excitotoxicity model in the nematode, the effect of central excitotoxicity modulators is conserved, including the neuroprotective effect of CREB/CRH-1.

**Figure 2:**
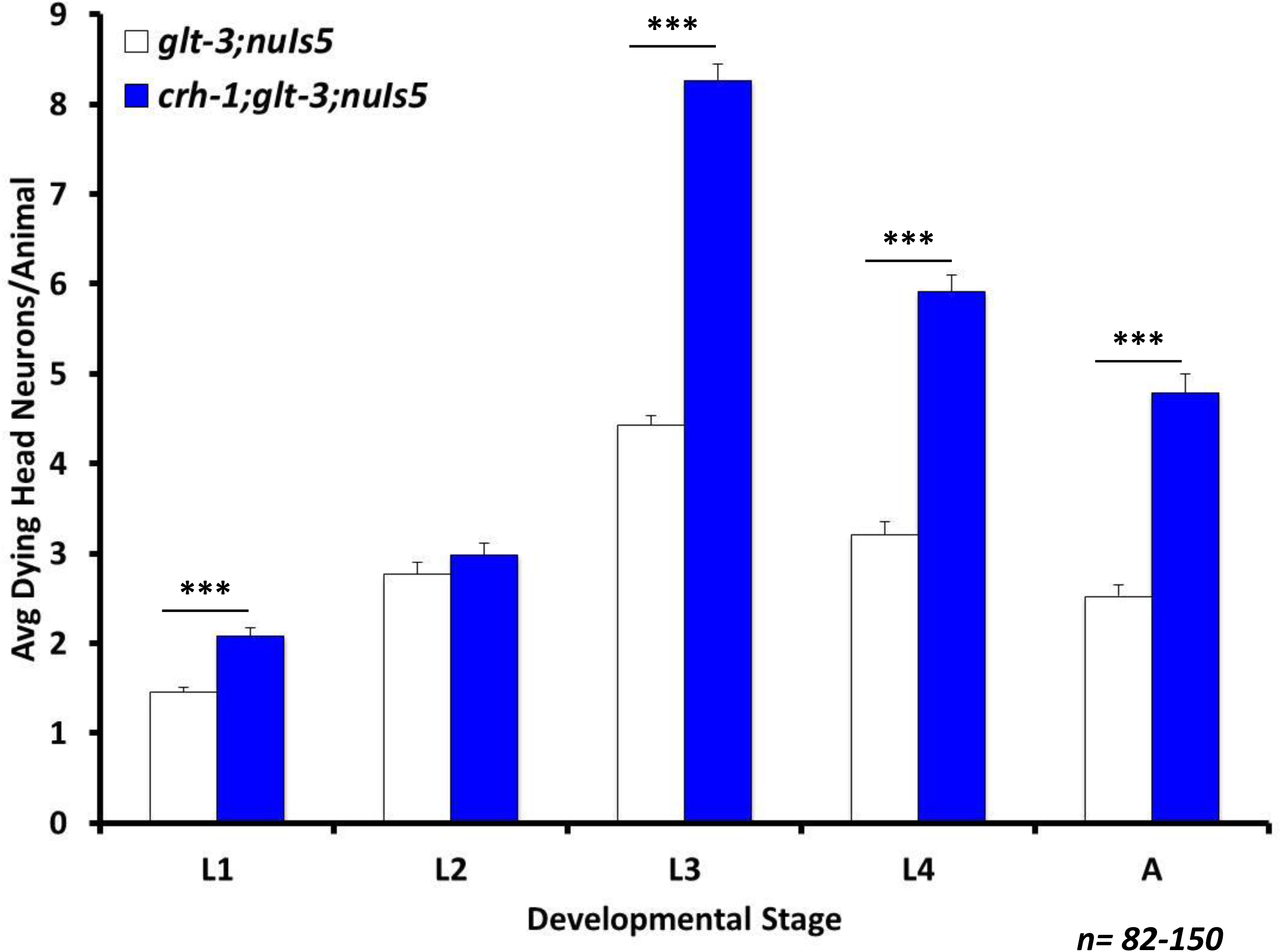
CREB/CRH-1 has a conserved role in neuroprotection Average number of degenerating head neurons per animal in different developmental stages, comparing the original excitotoxicity strain *(glt-3;nuIs5)* to a similar strain where CREB is eliminated *(crh-1;glt-3;nuIs5).* In this and all subsequent bar graphs: Error bars represent SE; Asterisks represent statistical significance of the difference between the indicated groups, where * indicates p<0.05; ** indicates p<0.01; *** indicates p<0.001

### The neuroprotective effect of CREB/CRH-1 is independent of neuroprotection by the Insulin/IGF-1 Signaling (IIS) cascade

Our previous studies on the regulation of excitotoxicity in nematodes and in mouse spinal cultures showed that excitotoxic neuroprotection is modulated by the Insulin/IGF-1 Signaling (IIS) cascade (Mojsilovic-Petrovic *et al.* 2009, Tehrani et al. 2014). Components of this cascade, and especially Akt, have also been suggested by others to mediate cross-talk with the CREB activation cascades (Soderling 1999, Walton & Dragunow 2000, Yuan & Yankner 2000). To determine if the neuroprotective effect of CREB is mediated by the same pathway as the IIS cascade we used genetic epistasis with a mutation in *age-1*, encoding the PI3 Kinase that is responsible for the activation of Akt. We have previously shown that the *age-1(hx546)II* mutation causes increased neuroprotection in nematode excitotoxicity (Mojsilovic-Petrovic et al. 2009). If *age-1’s* effect is mediated through CREB, then CREB/*crh-1 ko* should block the neuroprotective effect of the *age-1* mutation, and the *age-1;crh-1* combination should have the same neurodestructive effect as *crh-1 ko* alone. In contrast, if *age-1* works in a pathway that is independent CREB, then the combination of the neuroprotective *age-1* mutation and the neurodestructive *crh-1* mutation should result in an intermediate level of neurodegeneration. Our data shows (Figure 3) that the combination strain *age-1;crh-1;glt-3;nuIs5* has an intermediate level of neurodegeneration (between those seen for *age-1;glt-3;nuIs5* and *crh-1;glt-3;nuIs5).* We obtained a similar result using the PI3-kinase chemical inhibitor LY294002 in *crh-1;glt-3;nuIs5* animals (not shown). These results demonstrate that IIS and CREB consist of two independent signaling cascades that separately modulate necrotic neurodegeneration in nematode excitotoxicity. We therefore wanted to learn more on CREB’s mechanism of action.

**Figure 3:**
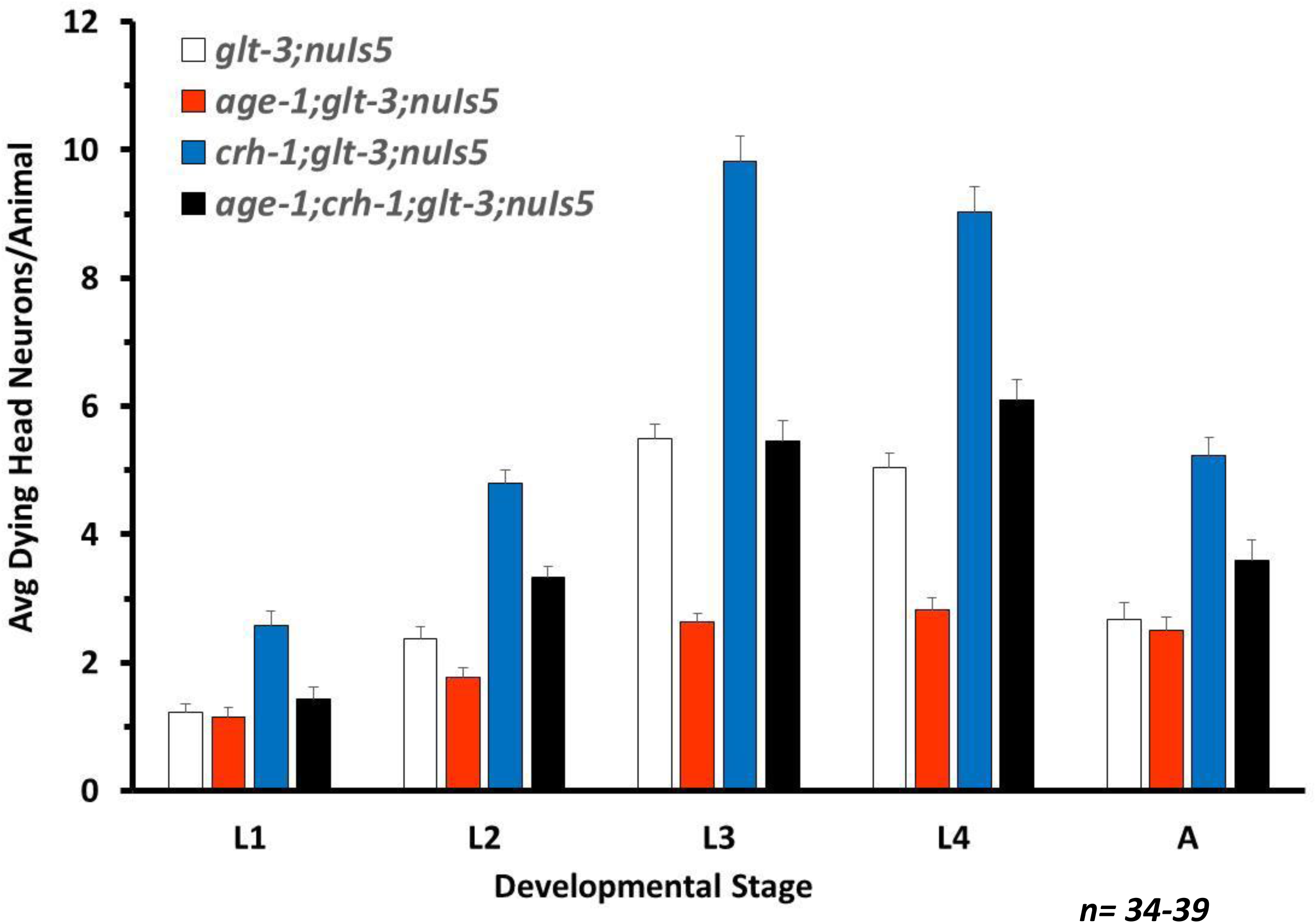
CREB/CRH-1 and the Insulin/IIS pathway regulate neuroprotection in separate pathways. Average number of degenerating head neurons per animal in different developmental stages, comparing the original excitotoxicity strain *(glt-3;nuIs5)*, CREB KO *(crh-1;glt-3;nuIs5)*, IIS neuroprotective mutation *(age-1;glt-3;nuIs5)*, and combination of CREB KO and IIS modification *(age-1;crh-1;glt-3;nuIs5)*

### The neuroprotection provided by CREB/CRH-1 in nematode excitotoxicity is more apparent in interneurons

To gain further insight into the process of neurodegeneration and neuroprotection we looked in more detail at the dying neurons and tried to identify them. Previously, our initial evaluation suggested that the overall pattern of necrosis of the at-risk neurons in the parental excitotoxicity strain *(glt-3;nuIs5)* is generally stochastic, so that each animal presents a different constellation of dying neurons (based on the rough location of the vacuole-like structures). However, more recently we suspected that the effect of CREB/CRH-1 might be preferentially pronounced in a specific subset of these at-risk neurons, which requires a more detailed analysis. The recognition of the dying neurons by name requires a careful examination of both the location & morphology of the neurons, and the characteristic structure of their processes. We take advantage of the GFP fluorescence of at-risk neurons in the excitotoxicity strain, and compare neuron location and the structure of its processes to the documented features of *glr-1* – expressing neurons (Brockie *et al.* 2001a, Brockie & Maricq 2006, Altun et al. 2002-2018). Usually, the GFP signal partially persists in the neurons as they swell-up and go through degeneration, and disappears only in the later stages of demise. We therefore used the persistent GFP label, as well as recording non-labeled neurons, to analyze the identity of degenerating neurons in a representative group of L3 animals.

We found that both in the regular excitotoxicity strain *(glt-3;nuIs5)* and in the excitotoxicity plus CREB-KO strain *(crh-1;glt-3;nuIs5)* the vast majority of swollen degenerating neurons are labeled, if weakly, with GFP. Very few swollen neurons do not show easily detectible GFP label. Of these non-green swollen neurons, most are found in locations that correlate with *glr-1* –expressing neurons that would normally be labeled with GFP (e.g., the RIG neurons, just posterior to the pharynx’s terminal bulb). Therefore, at least most of the non-green swollen neurons are likely to represent at-risk *(glr-1* –expressing) neurons where the GFP has already degraded. Excitotoxic neurodegeneration in our model, therefore, seems cell-autonomous. We noticed, however, that even though there is great variability in the probability of each neuron to die, there are also overall patterns that can be distinguished (Figure 4). While the large variability in degeneration levels prevents us from making conclusions on specific neurons (Figure 4A), combining the neurons into groups (Figure 4B) provides sufficient basis for determining the difference in degeneration-levels to be statistically significant. Under normal excitotoxic conditions *(glt-3;nuIs5)*, most of the neurons that die fall into the categories of the sensory neurons of the URY group, and the motor neurons of the RMD and SMD groups. In this original *glt-3;nuIs5* excitotoxicity strain, command interneurons such as AIB, AVA, AVB, AVD, and AVE die at lower frequencies. We further observe that the effect of CREB KO (in fold, comparing neurodegeneration levels with and without CREB) is particularly pronounced in the command interneurons. This difference in vulnerability can be based on a difference in the neuron’s exposure to the excitotoxic insult (e.g., difference in extracellular Glu concentrations, or a difference in the levels of expression of GluRs), or based on a difference in resilience of the postsynaptic neuron to a given insult. Since the command interneurons (which die less frequently) are reported to have particularly high levels of GluRs (Brockie et al. 2001a), we considered other mechanisms for differential vulnerability. To gain more insight into the mechanism of resilience, we study the pathway that allows CREB/CRH-1 to confer neuroprotection.

**Figure 4:**
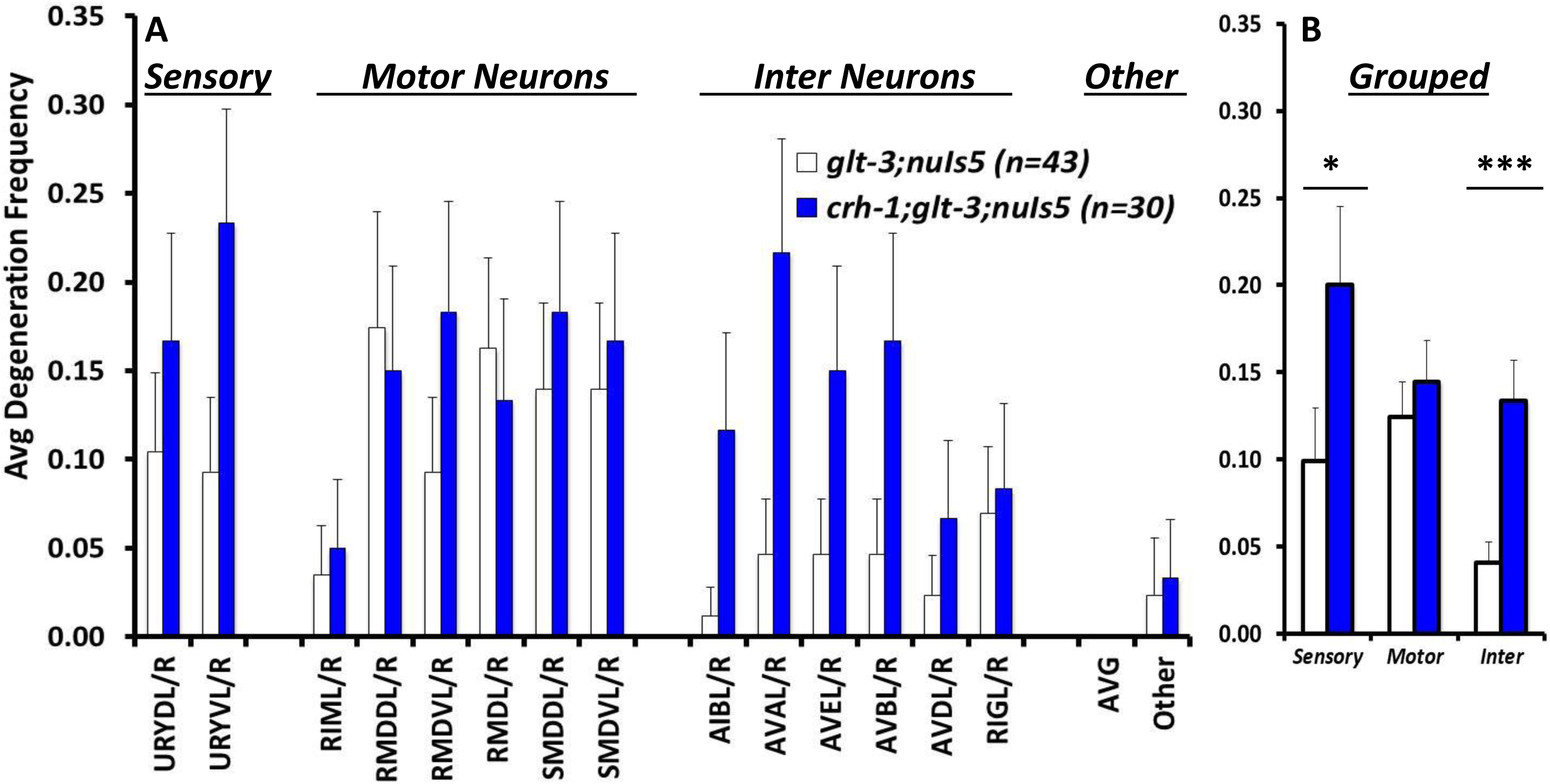
Neurodegeneration shows variability, possible patterns. A) Frequency of degeneration of identified neurons in the original excitotoxicity strain *(glt-3;nuIs5)* to a similar strain where CREB is eliminated *(crh-1;glt-3;nuIs5).* B) Pooled data from A, combining data for sensory neurons, motor neurons, and interneurons.

### Key components of the canonical cascade for CREB/CRH-1 activation do not contribute significantly to neuroprotection in nematode excitotoxicity

We next asked if the canonical mode of CREB activation by phosphorylation is involved in nematode excitotoxicity. Surprisingly, we have recently found that the CaMKK homolog CKK-1 (Eto *et al.* 1999), which is needed for a large potentiation of phospho-CREB –mediated transcription (Kimura et al. 2002, Yu et al. 2014, Moss et al. 2016), has no effect in nematode excitotoxicity (using the *ckk-1(ok1033) III* allele in the *glt-3;nuIs5* background) (Del Rosario et al. 2015). The effect of CaMK-IV in the nucleus is harder to decipher in the nematode, since the functions of both the cytoplasmic CaMK-I and the nuclear CaMK-IV are mediated in the nematode by a single nematode homolog, CMK-1 (Yu et al. 2014, Schild *et al.* 2014), which is expressed throughout the nervous system. The function of phosphorylation-dependent CREB partner CBP (which is also widely expressed in neurons (Hunt-Newbury *et al.* 2007)) can be studied in the worm using a *gain-of-function* allele of the *cbp-1* gene *(cbp-1(ku258)* III), encoding a CBP with a seven-fold increase in HAT activity (Eastburn & Han 2005). Similarly to the lack of effect of CaMKK/CKK-1, we find that extensive elevation of CBP/CBP-1 activity has no effect in nematode excitotoxicity (Figure 5). The lack of effect of *ckk-1* and *cbp-1* argues strongly against the involvement of canonical CREB activation in the neuroprotective effect of CREB in nematode excitotoxicity.

**Figure 5:**
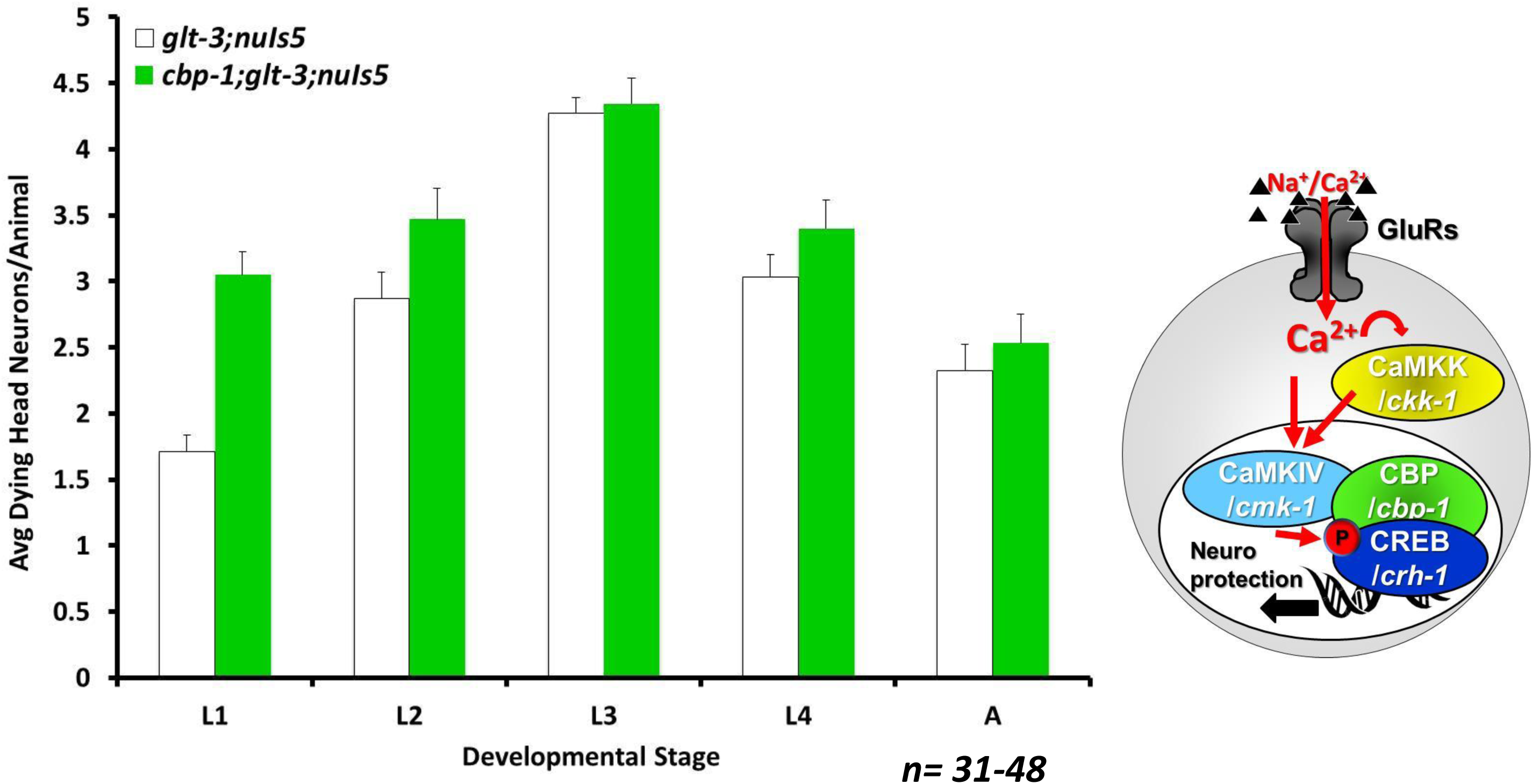
Canonical activator, CBP-1, has no effect on excitotoxic necrosis. Average number of degenerating head neurons per animal in different developmental stages, comparing the original excitotoxicity strain *(glt-3;nuIs5)* to a similar strain carrying also a hyperactivating mutation in CBP *(cbp-1;glt-3;nuIs5).*

### The non-canonical mediators of CREB/CRH-1 activation are central to neuroprotection in nematode excitotoxicity

Examining non-canonical mechanisms for CREB involvement in nematode excitotoxicity, we turned to test the CaMK-I/SIK/CRTC axis (Figure 6A). As mentioned above, the function of both CaMK-I and CaMK-IV is mediated in the worm by a single gene, *cmk-1.* In both canonical and non-canonical modes of CREB activation, the effect of WT CMK-1 is predicted to potentiate neuroprotection (either by directly activating CREB, as seen in Figure 1A, or by suppressing the inhibitor of it co-factor, as seen in Figure 1B/6A). Indeed, elimination of CMK-1 (using *cmk-1(ok287) IV* in the excitotoxicity strain) enhances neurodegeneration (equivalent to suppressing neuroprotection) (Figure 6B). Since this result cannot differentiate between the two modes of CREB activation, we advanced further down the pathway and examined the role of the SIK1/2 homolog AAK-2. We found that introducing the *aak-2(ok524)X* mutation to the excitotoxicity strain causes a pronounced decrease in neurodegeneration (equivalent to increasing neuroprotection) (Figure 6C). This observation is in line with a role for WT AAK-2 as a potentiator of excitotoxicity, and in line with the non-canonical mechanism of CREB activation. We finally checked the effect of the defining component of the non-canonical mechanism of CREB activation, CRTC. Elimination of CRTC (using the *crtc-1(tm2869)I* allele) had an extensive enhancing effect on the level of neurodegeneration, similar to the effect of eliminating CREB/CRH-1 itself (Figure 6D). Put together, these results suggest that the components of the non-canonical pathway for CREB activation are involved in neuroprotection from excitotoxicity. However, these results, in-and-of-themselves, do not provide information on whether they work together in a combined pathway, and on the hierarchical organization of the cascade.

**Figure 6:**
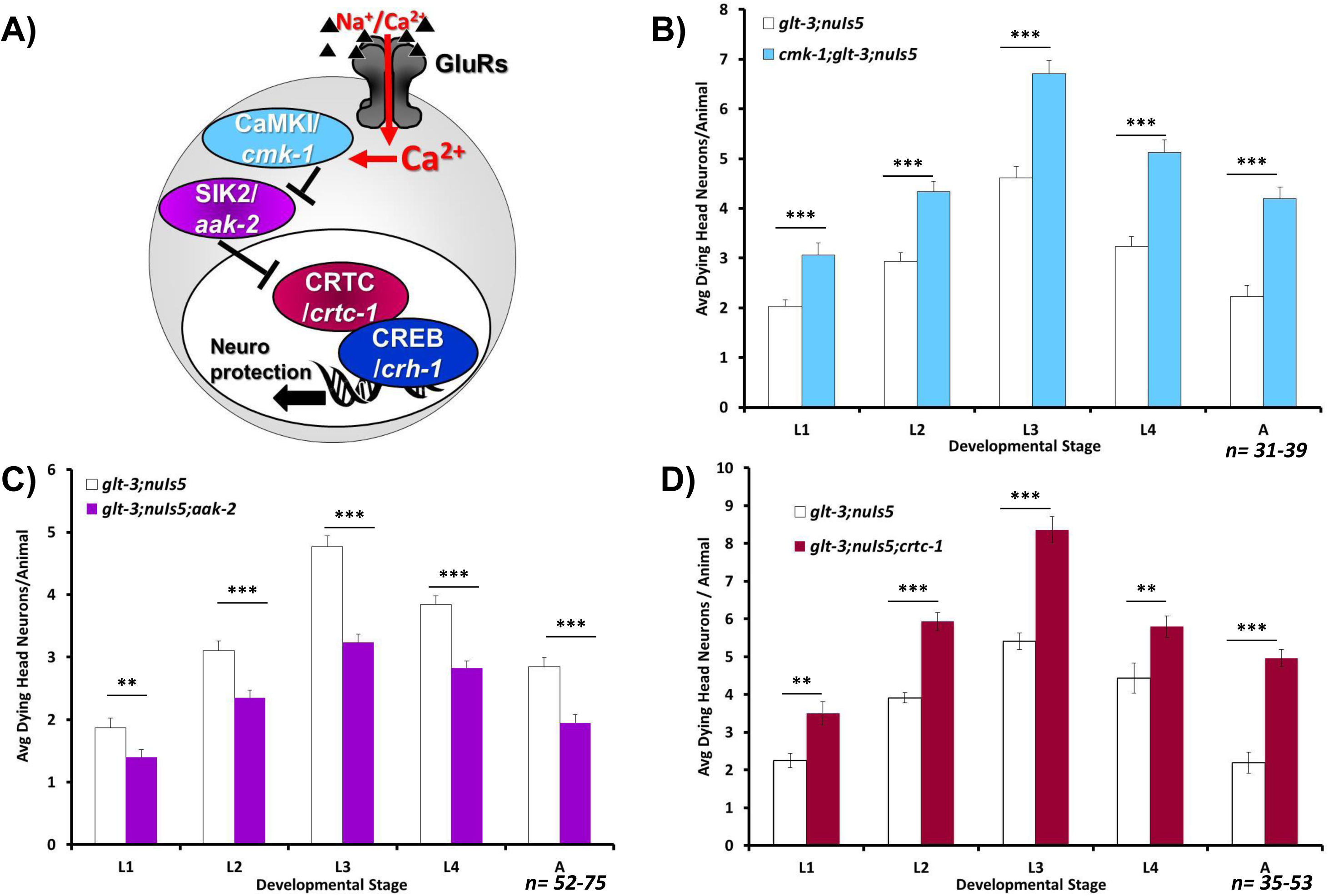
Non-canonical CREB modulators affect excitotoxic necrosis. Average number of degenerating head neurons per animal in different developmental stages, comparing the original excitotoxicity strain *(glt-3;nuIs5)* to a similar strain where a mediator of the non canonical pathway is eliminated: B) the CaMK-I/IV CMK-1; C) The SIK homolog AAK-2; D) CREB cofactor CRTC (CRTC-1).

### CRTC/CRTC-1 acts downstream of SIK2/AAK-2 to regulate neuroprotection in nematode excitotoxicity

AAK-2 has been previously shown to phosphorylate CRTC-1, thus determining its cytoplasmic vs nuclear distribution (Mair et al. 2011, Sasaki et al. 2011, Burkewitz *et al.* 2014, Altarejos & Montminy 2011, Burkewitz et al. 2015, Escoubas *et al.* 2017). However, AAK-2 has also been shown to regulate the activity of FoxO/DAF-16 to modulate levels of neurodegeneration (Greer *et al.* 2007, Tullet *et al.* 2014, Vazquez-Manrique *et al.* 2016). Since we have previously demonstrated that the IIS cascade and FoxO/DAF-16 regulates neuroprotection in nematode excitotoxicity (Mojsilovic-Petrovic et al. 2009, Tehrani et al. 2014), it becomes unclear if the effect of AAK-2 seen here is mediated by the CRTC-1/CRH-1 CREB cascade. We therefore used epistasis to test if SIK/AAK-2 and CRTC-1 act in the same pathway to regulate neuroprotection, by combining both the *aak-2* mutation and the *crtc-1* mutation in the excitotoxicity strain (Figure 7). Our results show that the neurodestructive effect of *crtc-1 ko* on nematode excitotoxicity remains intact in the absence of *aak-2* (whereas usually *aak-2 ko* is neuroprotective). These observations indicate that AAK-2 works in the same non-canonical CREB activation pathway as CRTC-1, and that CRTC-1 acts downstream of AAK-2, as suggested before for other effects of the AAK-2/CRTC-1/CRH-1 axis (Mair et al. 2011). We next wanted to see if other aspects of CRTC-1’s function are also in effect in nematode excitotoxicity.

**Figure 7:**
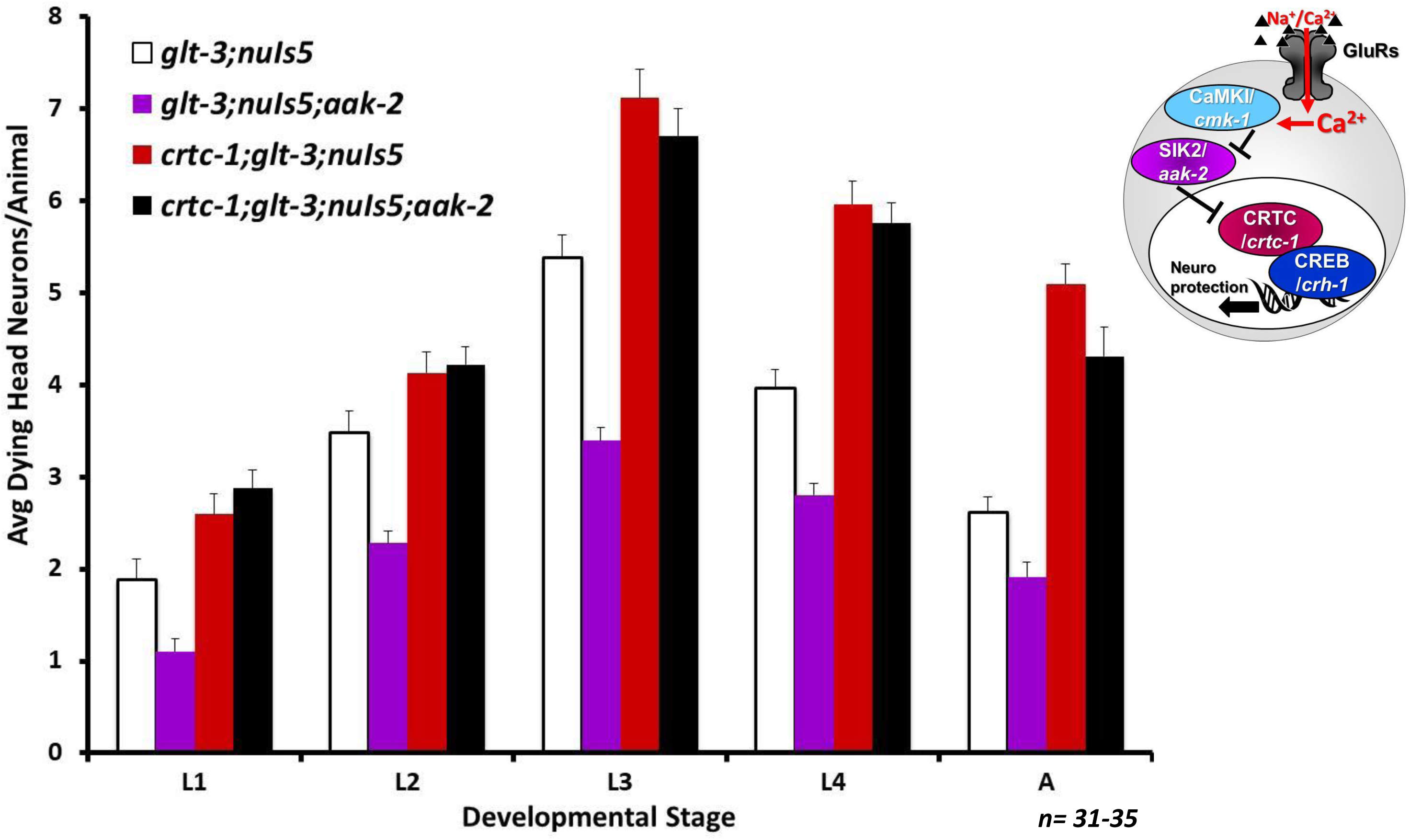
SIK/AAK-2 and CRTC-1 work in the same pathways. Average number of degenerating head neurons per animal in different developmental stages, comparing the original excitotoxicity strain *(glt-3;nuIs5)* to similar strain where SIK/AAK-2 is eliminated *(glt-3;nuIs5;aak-2)*, or CRTC-1 is eliminated *(crtc-1;glt-3;nuIs5)*, or both AAK-2 and CRTC-1 are eliminated *(crtc-1;glt-3;nuIs5;aak-2).*

### Our data is in line with the model of phosphorylation-dependent translocation of CRTC-1 between the cytoplasm and the nucleus in nematode excitotoxicity

Another prediction of the proposal that CRTC-1 is responsible for CREB activation in nematode excitotoxicity is that the hyperactivation of postsynaptic neurons causes CRTC to translocate into the nucleus of the at-risk neurons to mediate their protection, in accordance with previously suggested models (Mair et al. 2011, Burkewitz et al. 2015). Trying to gain visual access to this process we take advantage of a CRTC-1 labeled with red fluorescence developed by Mair & Dillin (Mair et al. 2011) and used it to monitor its translocation into the nucleus. We used a chromosomally-integrated transgene of *CRTC-1::RFP* expressed from its native promoter, and combined it with our excitotoxicity strain, where at-risk neurons are labeled with *P_glr-1_::GFP* (Figure 8). We observe that CRTC-1 is widely expressed in many neurons, including in *glr-1* expressing neurons. However, the expression was very dim (Figure 8 depicts highly contrasted images). Together with the small size of the head neurons, it was difficult to determine cytoplasmic vs nuclear localization in these neurons in a reliable and reproducible manner. Therefore, the limited quality of fluorescent labeling in these small neurons only allows us to deduce that CRTC-1 is available in the neurons where excitotoxicity takes place in our model.

**Figure 8:**
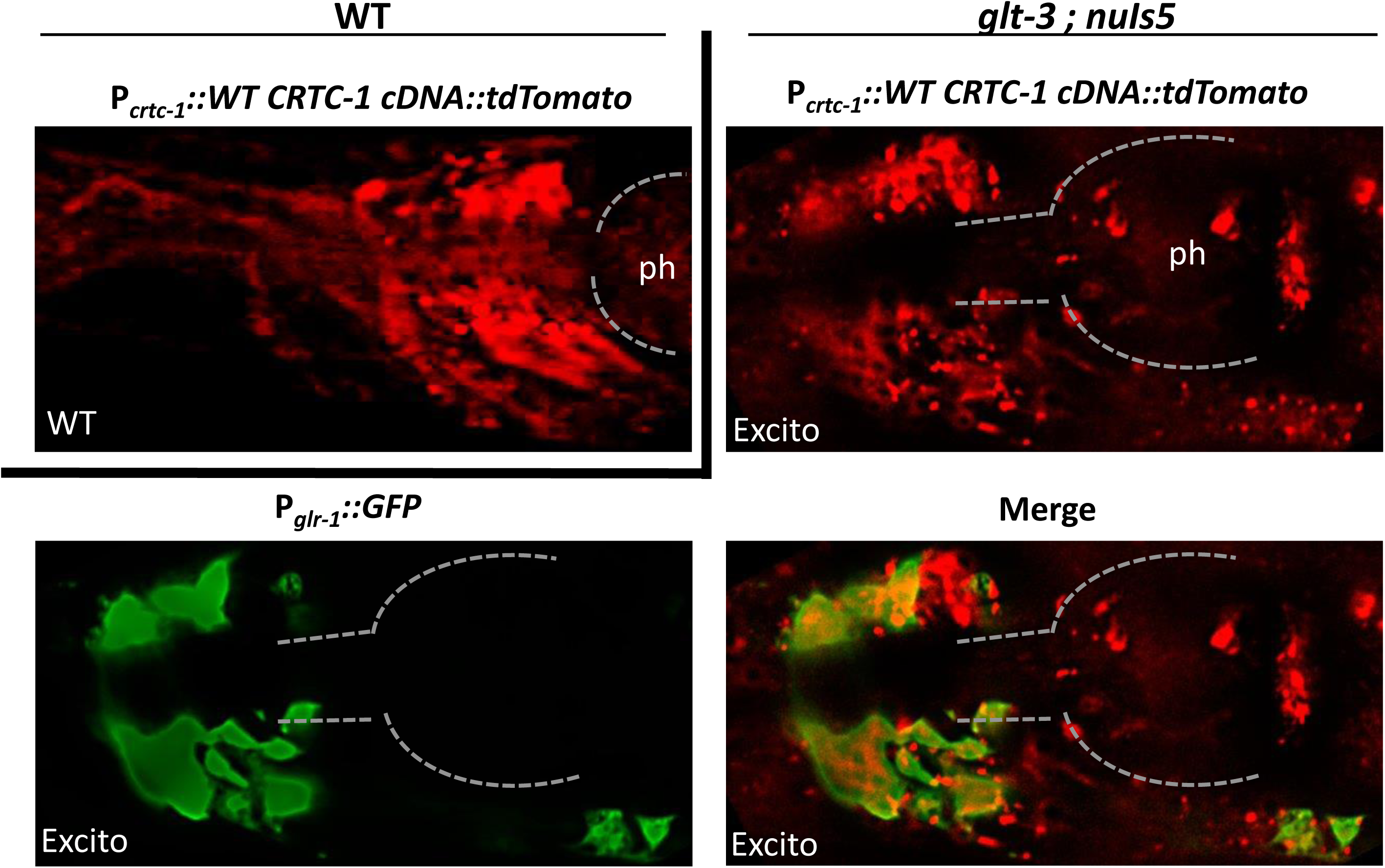
CRTC-1 is expressed in neurons exposed to insult. Epifluorescence images of collapsed z-stacks showing expression of WT CRTC in head neurons (red) compared to the location of GLR-1 expressing neurons (green). WT and excitotoxicity backgrounds are compared. Original red signal was very dim, and it was greatly enhanced by increasing contrast. The image shows the head area of representative animals (WT or excitotoxicity background). Dorsal is up, anterior is left, dashed line and the label “ph” mark the approximate area of the pharynx.

We further asked if the enhanced nuclear translocation reported previously for unphosphorylatable CRTC-1 (Mair et al. 2011) increases neuroprotection in nematode excitotoxicity. We used the Mair & Dillin unphosphorylatable mutant of CRTC-1, also fused to RFP and expressed from its native promoter *(CRTC-1(S76A,S179A)::RFP).* However, unlike the WT CRTC-1 transgene, this transgene is expressed as a non-integrated extrachromosomal array, a formulation that causes the transgene to be present at random in only some of the cells of the animal. This means that at any given animal only some of the neurons contain this transgene and can express the hyperactive non-phosphorylatable CRTC-1 it encodes. Our attempts to integrate this construct to stabilize expression were unsuccessful. We therefore tested the effect of suppressing neurodegeneration in our excitotoxicity strain with either the integrated transgene encoding WT CRTC expressed in *all* neurons or the non-integrated unphosphorylatable CRTC-1 expressed at random in only *some* neurons. We observe that the two transgenes reduced neurodegeneration to similar levels (Figure 9). The observation that the hyperactive unphosphorylatable CRTC-1 mutant is able to reduce neurodegeneration to the same level as WT CRTC-1 while being expressed in only a random subset of the affected neurons, is in line with a greater capacity of the mutant CRTC-1 to provide neuroprotection (though it cannot serve as a direct and independent proof for this effect).

**Figure 9:**
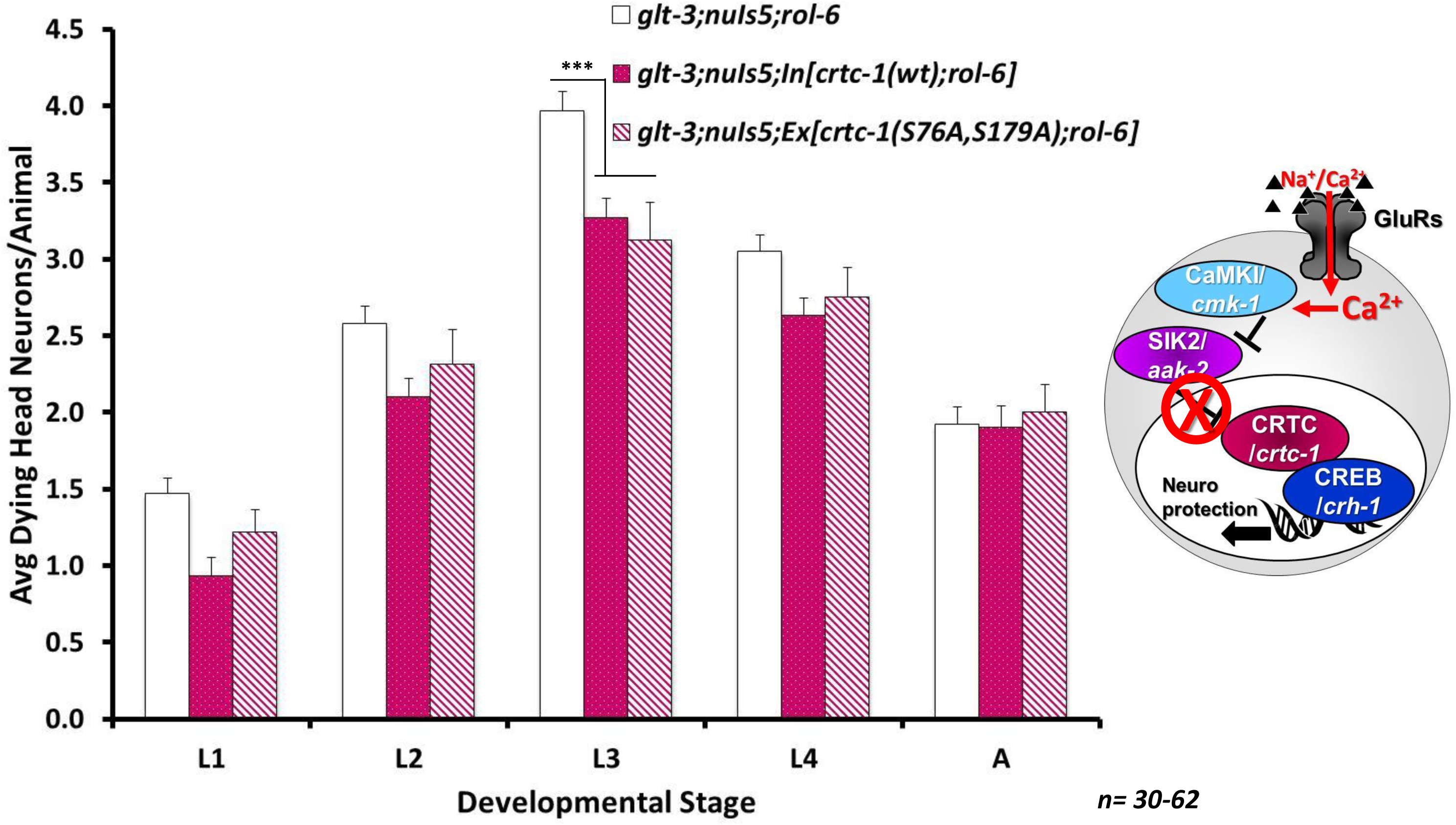
Unphosphorylatable CRTC-1 protects from excitotoxicity. Average number of degenerating head neurons per animal in different developmental stages, comparing the original excitotoxicity strain *(glt-3;nuIs5)* to a similar strain carrying also an integrated transgenic construct expressing cDNAs encoding WT CRTC-1 or an extrachromosomal nonintegrated construct expressing S76A,S179A non-phosphorylatable CRTC-1.

### CREB/CRH-1 activity in nematode excitotoxicity is cell autonomous and independent of its phosphorylation

To firmly establish that the neuroprotection provided by CREB in nematode excitotoxicity is turned on via the phosphorylation-independent, non-canonical pathway, we used a phosphorylation-deficient mutant version of CREB (Kimura et al. 2002). The conserved site of phosphorylation in the transactivation / KID domain of nematode CREB/CRH-1 has been determined to be Ser29, and previous studies showed that an S29A mutant construct was unable to give rise to CaMKIV/CMK-1 or CaMKK/CKK-1 - induced transcription by CRH-1. We therefore compared the ability of WT and S29A mutant *crh-1* cDNA to rescue the effect of *crh-1 ko* and reduce neurodegeneration towards more normal levels. To address the question of cell autonomous/non-autonomous effect of CREB in our model, we expressed the WT and mutant *crh-1* cDNA under the *glr-1* promoter (i.e., in the at-risk neurons). Both transgenes were expressed as non-integrated extrachromosomal constructs. We observe that both WT and S29A non-phosphorylatable CREB/CRH-1 can rescue the neurodegeneration (partially, as expected from a non-integrated construct), indicating a cell-autonomous effect. Importantly, the two versions of CREB/CRH-1 rescue neurodegeneration to the same extent (Figure 10). This observation indicates that CREB/CRH-1 does not need to be phosphorylated (at least not in the canonical phosphorylation site in its transactivation domain) to provide neuroprotection. Together, these results firmly establish that CREB-mediated neuroprotection in nematode excitotoxicity is achieved cell autonomously and is independent of CREB activation by canonical phosphorylation in its transactivation domain.

**Figure 10:**
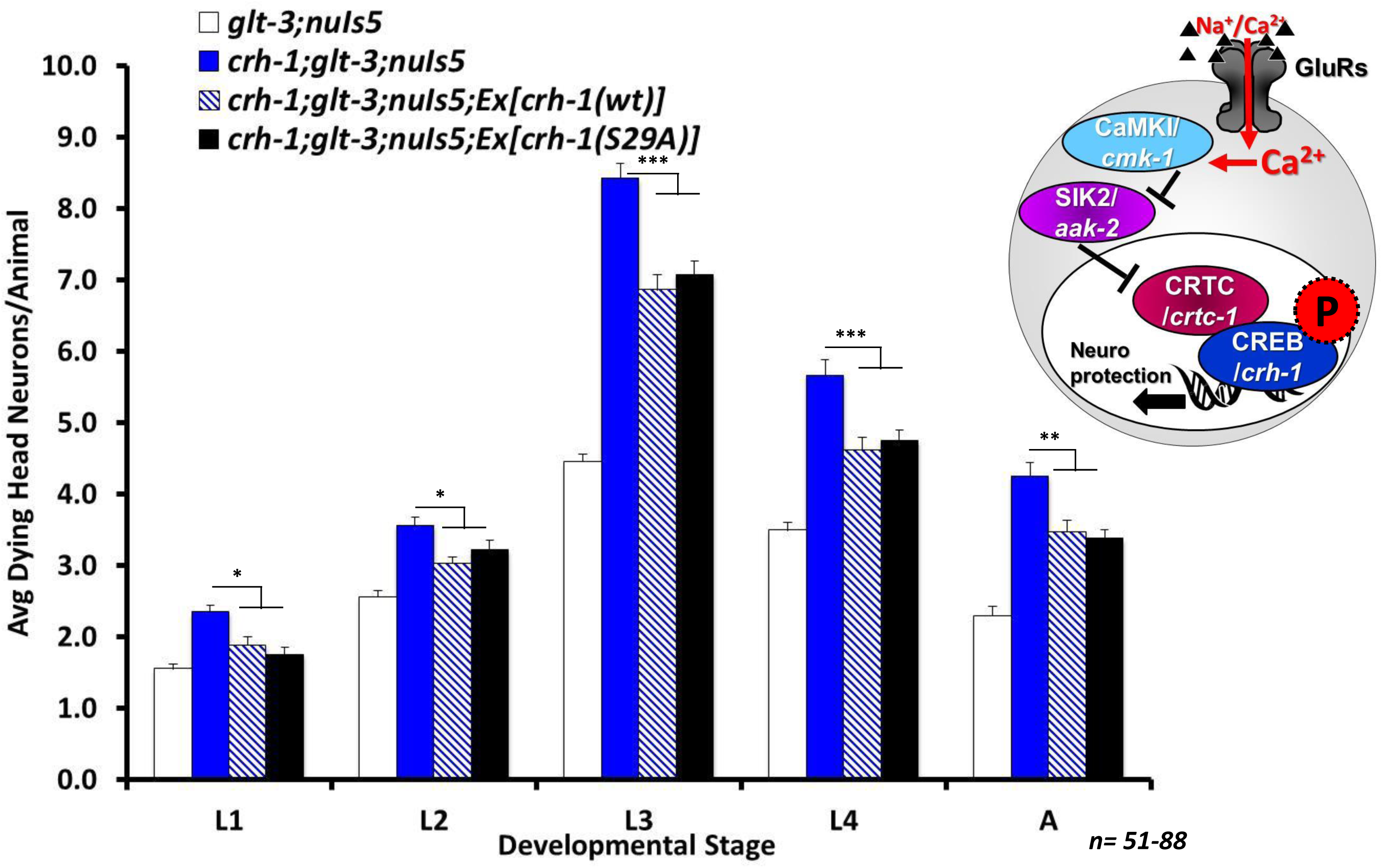
CREB’s ability to protect from excitotoxic necrosis is independent of its phosphorylation status. Average number of degenerating head neurons per animal in different developmental stages, comparing the original excitotoxicity strain *(glt-3;nuIs5)* to a similar strain carrying also extrachromosomal non-integrated transgenic constructs expressing cDNA encoding WT or a S29A mutant non-phosphorylatable CRH-1.

## Discussion

In this study we establish that the non-canonical mode of CREB activation mediates protection from necrotic neurodegeneration in nematode excitotoxicity. Our data shows that the neuroprotective function of CREB in excitotoxic necrosis (Figure 2) is conserved over a large evolutionary distance, suggesting that it forms a critical core biochemical cascade in neuroscience. We also show that the effect of the IIS cascade on neuroprotection is separate from the effect of CREB (Figure 3).

Unlike other forms of cell death in *C. elegans* (Lettre & Hengartner 2006), the pattern of cell death in excitotoxic necrosis is not stereotypic, showing considerable variability (Figure 4A). Nonetheless, it also shows a clear pattern of susceptibility and resilience: under basal excitotoxic conditions, most of the dying neurons are the URY sensory neurons and the RMD & SMD motorneurons; under these conditions, the command interneurons are mostly protected, and they degenerate in large numbers only when CREB/CRH-1 is eliminated (Figure 4B). It is not immediately clear what causes the special vulnerability of the RMD and SMD neurons to excitotoxic neurodegeneration in the WT background, and what is the source of CREB’s ability to protect the command interneurons. Generally, we try to differentiate between susceptibility that is caused in some neurons because they are exposed to a greater Glu insult, versus resilience that comes from more effective protection from a similar insult. The resilience of the command interneurons (when CREB is present) is in stark contrast to the fact that they express the highest levels of all GluRs (Brockie et al. 2001a), arguing against susceptibility that is based at the level of the GluR. Examining resilience, we note that CREB (and its activating factors) is reported to be expressed in low levels in many neurons (Kimura et al. 2002), while there’s high expression in very few neurons (Chen et al. 2016) does not correlate with the neurons studied here. Indeed, a recent study suggests that CREB is active in *glr-1* –expressing neurons (Moss et al. 2016) (although this study suggests that active CREB/CRH-1 moderately reduces *glr-1* transcription, while we observe that *crh-1 ko* affects mobility in the opposite direction, Supplementary Figure 1). Therefore, at this point, there is no support for a model of susceptibility and resilience that is based of *postsynaptic* properties, such as increase GluR expression, or reduced expression of CREB and its activating cascade. A possible *presynaptic* explanation to the differential susceptibility is that the sensitive neurons receive more/stronger synaptic inputs. In *C. elegans*, the number of synaptic inputs (as counted in EM reconstruction of the nervous system) is suggested to serve as an indication on the strength of signaling between these neurons (Gray *et al.* 2005, Leinwand & Chalasani 2013). We therefore analyzed publicly available data on the number of synapses that each of these neurons receives (White et al. 1986, Jarrell *et al.* 2012) (using data from WormWeb.org & WormWiring.org), and used the map of glutamatergic neurons (Serrano-Saiz *et al.* 2013) to identify which of these connections is glutamatergic. Tallying the number of glutamatergic inputs that each of the *glr-1 –* expressing neurons receives (Supplementary Table 1), we notice that there is no correlation between the number of incoming glutamatergic connections and the tendency of the neurons to die by excitotoxicity (see RMDs vs AVA, or SMDs vs AVB, AVD, and AVE). One observation that does correlate with the observed neuronal susceptibility is the location of these synapses in the nerve ring: the SMD and RMD synapses are located in the inner-most aspect of the nerve ring, close to the space between the pharyngeal muscle and the nerve ring (Supplementary Figure 2 and WormWiring.org), a compartment washed by body fluids (Altun et al. 2002-2018). In contrast, the inputs to the command interneurons are located further laterally in the radial organization of the nerve ring, and are less exposed to the centrally located body fluid compartment. URYs are also exposed to the body fluids. The buildup of Glu in our excitotoxicity model is specifically achieved by a KO of *glt-3*, a GluT that regulates Glu concentrations in the body fluids (Mano et al. 2007). Furthermore, our unpublished studies using GCaMP imaging to test the effect of different GluT KO on different synapses further support that *glt-3 ko* exerts its effect radially from the inner face of the nerve ring (Lee, Chan, and Mano, manuscript in preparation). It therefore seems that the neurons most susceptible to excitotoxic necrosis in our model are those whose synapses are most exposed to elevation in Glu concentrations caused by KO of *glt-3.* In contrast, neurons whose synapses are further away from the compartment of high Glu concentrations are protected, as long as CREB signaling is intact. It therefore seems that CREB/CRH-1 has the greatest neuroprotective capacity to make a difference in viability fate in a “penumbra-like” region of moderate Glu insult experienced by the command interneurons.

Further asked how CREB is activated in nematode excitotoxicity. We demonstrate that canonical mediators of CREB activation and function, such as CaMKK/CKK-1 and CBP/CBP-1 have little to no role in excitotoxic necrosis in the nematode (Figure 5 and our previous studies (Del Rosario et al. 2015)). We further demonstrate that the mediators of the non-canonical route have a very strong effect on CREB’s activation (Figure 6), and that the cascade is arranged in the same order (Figure 7) as the one suggested in mammals (Altarejos & Montminy 2011, Sasaki et al. 2011). Our data shows that neuroprotection hinges on CRTC-1, which is widely (though not intensely) expressed in the nematode’s nervous system (Figure 8). Though not conclusive on their own, our observations are in line with previous studies showing that CRTC-1 is regulated by phosphorylation to translocate to the nucleus in the affected neurons (Figure 9). Finally, our data clearly shows that CREB/CRH-1 acts cell autonomously and independently of its phosphorylation at the conserved site in the transactivation domain, to provide neuroprotection in excitotoxic necrosis in the nematode. These observations suggest that although the canonical route of CREB activation is in widespread use in many scenarios in both mammals and nematodes, the mode of CREB activation that is conserved in protection from excitotoxic necrosis, is the non-canonical one. This uncommon mode of activation might be involved in providing at least partial specificity, allowing CREB to turn on different sets of genes depending on the specific conditions in the cell.

The phosphorylation of CREB in the transactivation domain has been the cornerstone of suggested mechanisms of CREB activity for many years. It serves both as a crossroad for many signaling pathways, and, together with the phosphorylation-dependent binding of the histone acetyl transferase (HAT) CBP, as a main mechanism for transcriptional activation. A number of recent studies have shown that in some important cases the non-canonical pathway is at play, involving CREB stabilization on the DNA by CRTC (which has its own transactivation domain), and the potential recruitment of other HATs with other histone acetylation profiles. However, it remained unclear what is the balance of significance between the canonical and non-canonical models. The fact that the non-canonical model is the one conserved through evolution to provide neuroprotection in excitotoxic necrosis adds much emphasis to this mode of CREB activation in this specific scenario. The non-canonical mode of CREB activation, with its expected ability to give rise to histone acetylation patterns that are different from those of the canonical mode of activation (Uchida et al. 2017), might direct the differential expression of target genes that were not studied before. Our findings do not negate the significance of canonical CREB activation in excitotoxicity that leads to apoptosis. However, necrotic neurodegeneration, which is much less understood, is a leading mediator of neurodegeneration in brain ischemia (including in the non-immediate stages of neurodegeneration, when neurons are “stunned” and need to choose their survival fate). Finding the right handles that might tip the balance towards specific CREB activation, and identifying the most critical neuroprotective genes associated with this specific mode of CREB activity, will be crucial for the future development of therapeutic interventions in brain ischemia.

## Acknowledgments. Conflict of interest disclosure

We would like to thank the Li lab (especially A. Alexander and J.-S. Yang) and the Emerson lab for help with molecular biology and microinjections. We would like to thank all members of the Mano lab, A. Alexander, and A. Khan for technical support and critical reading of this manuscript. We would also like to thank M. Driscoll and H. Kuroyanagi for gift of plasmids. We thank the *Caenorhabditis* Genetic Center at the Univ. of Minnesota (which is funded by NIH Office of Research Infrastructure Programs, P40 OD010440), and the Japanese National Bioresource Project (NBRP, Ministry of Education, Culture, Science, Sports and Technology, Japan) for providing strains. This project was supported by The Alexandrine and Alexander L. Sinsheimer Fund (P60134 to IM), the American Heart Association (16GRNT31500004 to IM), and NIH/NINDS (NS096687 to IM). The Mano lab is also supported by NIH/NINDS (NS098350 to IM), and by an institutional RCMI grant (G12RR003060-26 to CCNY).

The authors declare no conflict of interest.

---

**Supplementary Figure 1:**
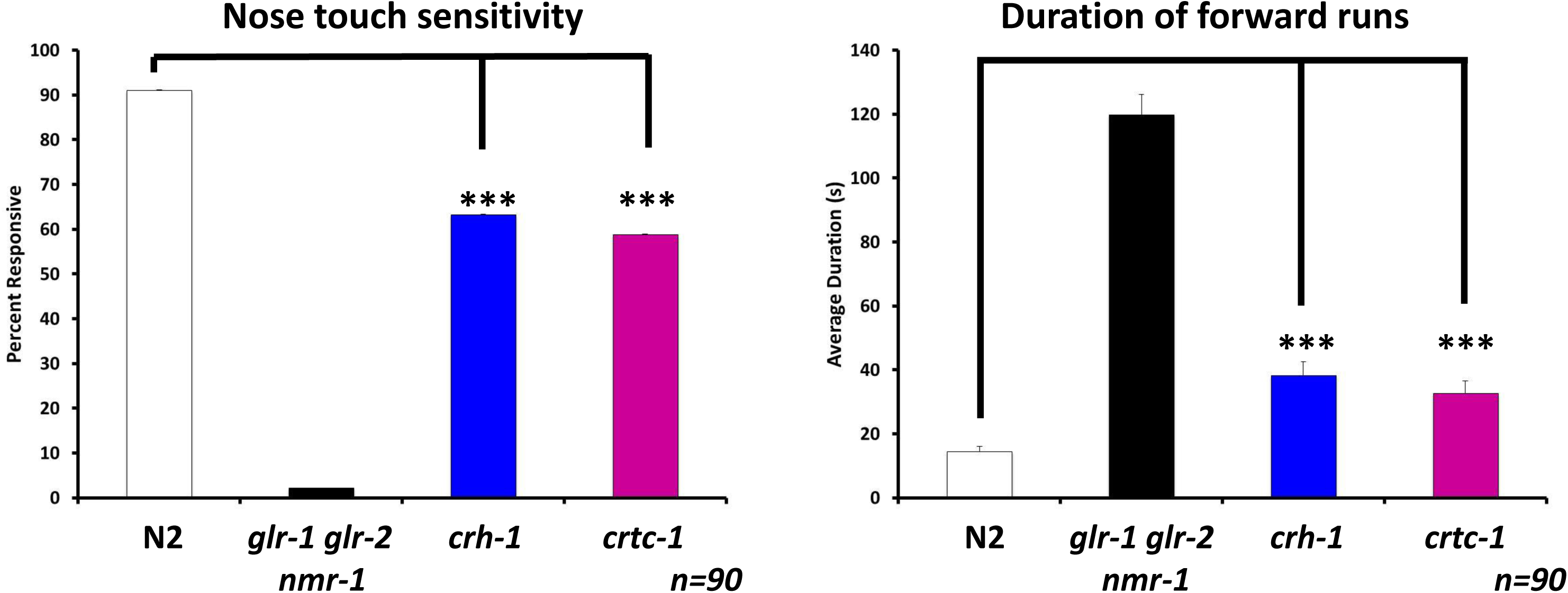
Behavioral tests of synaptic strength in Glu connections to command interneurons. Left: Nose touch sensitivity; Right: duration of spontaneous forward runs in WT, CREB, and CRTC mutants (with a strain without GluR as a point of comparison).

**Supplementary Figure 2:**
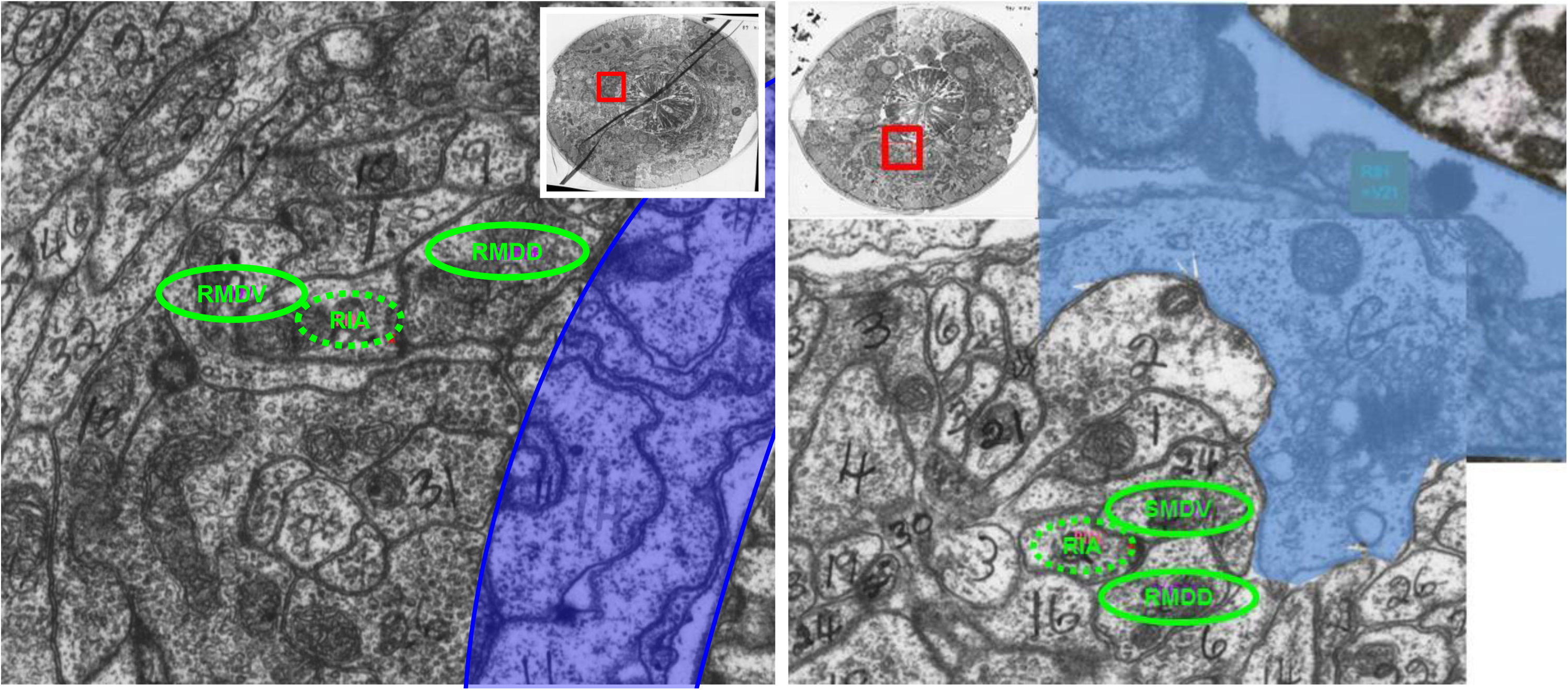
The synapses between RIA and the RMD/SMD neurons are located on the innermost aspect of the nerve ring. Images represent a coronal section through the nerve ring where specific synapses were identified, with insert showing the overall area. Images, cell identification and synapse location is taken (and modified-) from WormWiring.org. Light blue shaded area represent space between the pharynx and the neurons that is accessible to body fluids (limited by the basal membrane found at the innermost aspect of the nerve ring, see WormAtlas.com). Green ovals denote neuron identities, with dashed line indicating the presynaptic neuron and solid line representing the postsynaptic neuron.

**Supplementary Table 1:** Counting of glutamatergic inputs to *glr-1* expressing neurons (Synapse numbers based on WormAtlas.org, Wormweb.org/neuralnet, identification as glutamatergic based on Serrano-Saiz et al., Cell 155, 659–673, 2013)

| Neuron | Glutamatergic Presynaptic Cells (# of Synapses) | Total |
| --- | --- | --- |
|  | *Sensory Neurons* |  |
| URYs | IL1(2); OLL(1) | 3 |
|  | *Motor Neurons* |  |
| RMDs | ALM(2); IL1(56); OLL(26); OLQ(15); RIA(105); RIM(6); URY(34) | 244 |
| SMDs | ADA(4); AIB(8); AIZ(1); OLL(19); RIA(69); RIM(12); URY(14) | 127 |
|  | *Interneurons* |  |
| AVAs | ADA(3); ADL(7); AIB(3); AQR(5); ASH(7); AUA(4); AWC(1); BAG(4); DVC(12); FLP(48); LUA(21); PHB(29); PHC(1); PLM(5); PQR(19); PVD(27); RIM(3); URY(1) | 200 |
| AVEs | ADA(2); AIB(1); AIM(1); AIZ(11); ALM(1); ASH(3); AUA(7); BAG(9); FLP(6); IL1(4); OLL(37); RIG(5); URY(17) | 104 |
| AIBs | ADA(5); ADL(18); AIN(2); AIZ(21); ASE(23); ASG(5); ASH(8); ASK(2); AWC(7); BAG(1); DVC(4); FLP(5); RIM(4) | 105 |
| AVBs | ADA(18); ADL(7); AIB(8); AIM(2); AQR(7); ASH(9); AVM(12); DVC(1); FLP(15); PVQ(1); PVR(8); RIM(12); URY(1) | 101 |
| AVDs | ADA(1); ADL(12); AIM(3); ALM(1); AQR(2); ASH(12); AUA(1); FLP(30); LUA(10); PHA(1); PHB(3); PLM(5); PQR(13) | 94 |
| RIMs | ADA(8); AIB(32); AIZ(8); ASH(1) | 49 |
| RIGL/R | AIB(1); AIZ(1); BAG(10); DVC(10) | 22 |
|  | *Other* |  |
| AVG | PHA(8); PQR(1) | 9 |

